# Plus and minus ends of microtubule respond asymmetrically to kinesin binding by a long range directionally driven allosteric mechanism

**DOI:** 10.1101/2021.08.30.458176

**Authors:** Huong T. Vu, Zhechun Zhang, Riina Tehver, D. Thirumalai

## Abstract

Many members in the kinesin superfamily walk predominantly towards the plus end of the microtubule (MT) in a hand-over-hand manner. Despite great progress in elucidating the mechanism of stepping kinetics, the origin of stepping directionality is not fully understood. To provide quantitative insights into this important issue, we represent the structures of conventional kinesin (Kin1), MT, and the Kin1-MT complex using the elastic network model, and calculate the residue-dependent responses to a local perturbation in these constructs. Fluctuations in the residues in the *β* domain of the *α/β*-tubulin are distinct from the *α* domain. Surprisingly, the Kin1-induced asymmetry, which is more pronounced in *α/β*-tubulin in the plus end of MT than in the minus end, propagates spatially across multiple *α/β*-tubulin dimers. Kin1 binding expands the MT lattice by mechanical stresses, resulting in a transition in the cleft of *α/β* tubulin dimer between a closed (**CC** for closed cleft) state (not poised for Kin1 to bind) to an open (**OC** for open cleft) binding competent state. The long-range asymmetric responses in the MT, leading to the creation of **OC** states with high probability in several *α/β* dimers on the plus end of the bound Kin1, is needed for the motor to take multiple steps towards the plus end of the MT. Reciprocally, kinesin binding to the MT stiffens the residues in the MT binding region, induces correlations between switches I and II in the motor, and enhances fluctuations in ADP and the residues in the binding pocket. Our findings explain both the directionality of stepping and MT effects on a key step in the catalytic cycle of Kin1.

## Introduction

The motor protein, kinesin, utilizes chemical energy generated by ATP hydrolysis, to transport organelles and vesicles, and drives the motility of spindles and chromosomes inside the cell. To carry out these functions, it interacts with microtubule (MT), an important component of the cytoskeleton. For example, conventional kinesin, referred to as Kin1, walks on MT, assembled from tubulin dimer, towards the plus-end of MT [1, 2, 3, 4, 5], in the process transporting organelles from the nucleus towards the cell membrane. In contrast, kinesin-13, part of the kinetochore that tethers chromosome, depolymerizes the spindle MT to drive chromosome segregation during mitosis [6]. In much of the discussions of the kinesin stepping mechanism, it is tacitly assumed that MT may be treated as a passive polar track [7]. Indeed, not only does MT impact the stepping of the motors, but also other functions involving transportation [8, 9, 10, 11]. In particular, the kinesin-MT interaction sets the stepping direction and enhances rate-limiting ADP release by about three orders of magnitude [12]. The goal of the present study is to provide a structural explanation for the origin of directionality in the Kin1 stepping using the notion of local stiffness changes in the tubulin dimers and the motor, triggered by Kin1-MT interactions.

We were motivated to undertake this study based on the following observations: (1) It is known that kinesin-MT interaction affects the MT [13]. For instance, Kin1 alters the microtubule lattice upon binding [14, 15], thus establishing that the MT is not merely a static stiff polar track but plays a dynamic role in controlling the motility of the motor. (2) We showed previously that, after the tethered head detaches and stochastically diffuses towards the next binding site on the MT [7, 16], the Kin1-MT interaction poises the free head to bind with the correct orientation [7]. (3) Expansion of the MT lattice induced by Kin1 binding [10, 14, 15] and even removal of *α/β* dimers when the motor steps on MT [17] shows that the polar track must play a dynamic role.

Here, we use the Structural Perturbation Method (SPM) [18, 19], within the framework of Normal Mode Analysis of the kinesin-MT complex [20, 21, 22], to calculate the changes in the residue fluctuations in kinesin and MT. The SPM calculations are used to address the following questions: (i) Are there residue dependent responses in kinesin upon MT binding and vice versa, and how can they be quantified? (ii) Is the response of *αβ*-tubulin at the plus and minus ends of the MT distinct? (iii) Why does motor binding at a specific site on the MT promote other kinesin molecules to bind to the same protofilament [23, 14]? (iv) Does the kinesin induced expansion of the MT lattice [14, 15] promote binding of other kinesins preferentially towards the plus end of the MT?

We answer these questions by first piecing together different PDB structures for kinesin and MT to create kinesin-MT complexes. Next, we created elastic network models (ENM) in which contacts present within a cut-off distance in the complexes are assumed to interact via harmonic potentials. The eigenvalues and eigenvectors of the ENM (see Methods) and how they change upon perturbation are used to assess the residue-dependent dynamic responses throughout the different complexes, thus allowing us to answer the questions posed above. Binding of Kin1 to the MT produces an asymmetric response with residues in the *β*-tubulin exhibiting larger fluctuations than in the *α*-tubulin. Surprisingly, the asymmetric response, which is more prominent in the plus end tubulin dimers, propagates spatially across the MT protofilament. Kin1 binding at the cleft of the intra-dimer interface of a specific *α/β* results in a transition from a closed cleft (**CC**) to an open cleft (**OC**) state. The mechanical stress created in this process poises the downstream (plus end) *α/β* dimers to be in the **OC** states, which enhances the probability of the motor stepping towards the plus end, thus minimizing backward steps. As a corollary, it follows that upon binding of a single Kin1 other motors would preferentially bind to the sites on the plus end of the MT.

## Results and Discussion

### Kin1 binding elicits an asymmetric response between the plus and minus end tubulins

In an interesting experiment [24], it was observed that binding of Kin1 to MT is cooperative. Relative to a stationary Kin1, (achieved by coating Kin1 by fluorescent beads), which hydrolyzes ATP without stepping, a second Kin1 is more likely to bind to the plus-end of the same MT [24]. The effect of cooperative binding propagates up to distances on the order of a ~ few microns. This finding is surprising because a single MT protofilament is essentially one dimensional, and all the interactions are short-ranged, which implies that long range correlations are unexpected.

The experimental observations [24] raise the possibility that Kin1 binding to MT affects the fluctuations of the residues at the plus-end differently than the residues at the minus-end. To test this hypothesis, we constructed a protofilament consisting of 4 potential kinesin binding sites. We first calculated the amplitude of fluctuations of each MT residue using a structure without a bound Kin1. Next, we created a complex with a Kin1 dimer attached to the two middle binding sites in the MT segment, leaving a pair of *α/β* tubulins free at the plus and minus ends (referred to as (Short)_1_-DU-Near complex, see Figure 1a). We then calculated the change in tubulin fluctuations at the minus-end, 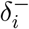, and the plus-end, 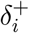 (Methods). To determine whether there is an asymmetry in the response of a tubulin, we calculated 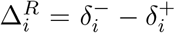 (Equation 9). A non-zero value of Δ^*R*^ would imply that the two ends of the tubulin are affected differently upon Kin1 binding.

**Figure 1:**
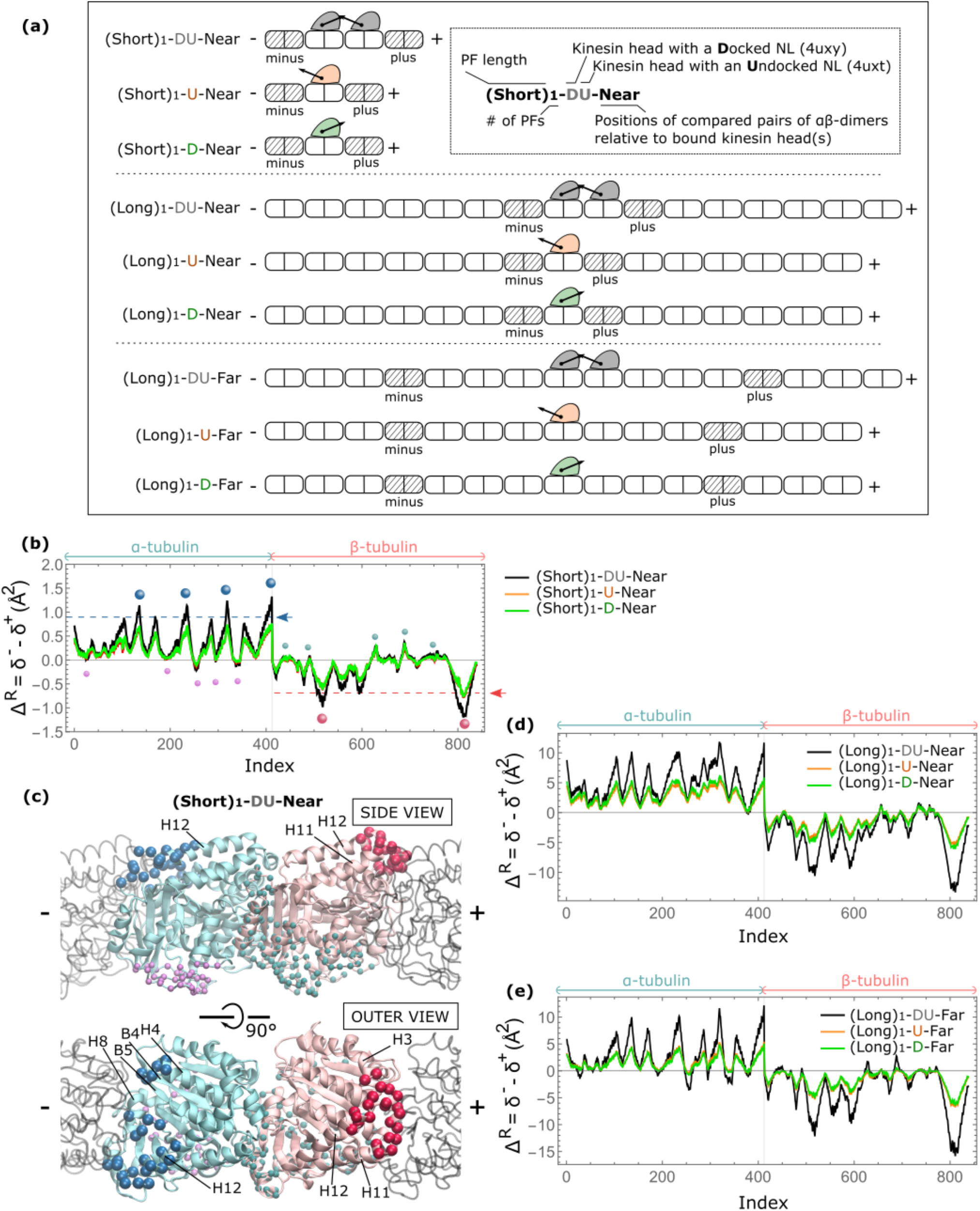
Asymmetric response of the plus and minus ends of tubulin upon Kin1 binding. (a) Schemes of different constructs used in this figure. Each construct has one or two Kin1 motor domains (filled objects) bound to a short or long single protofilament consisting of multiple *αβ* tubulin dimers (repeated opened objects). The docked (undocked) neck linker is represented as arrows pointing towards the plus (minus) ends of the protofilaments. The minus and plus *αβ* tubulin dimers used to calculate the asymmetric response are highlighted with stripes. On the left are labels for each construct. Inset shows the meaning of the labels. (b) Difference in the responses of *αβ*-tubulin residues upon Kin1 binding, Δ^*R*^, calculated for the constructs with single extra dimer at plus and minus ends where Kin1 motor domain binds (short protofilaments). Residues belong to *α* and *β* tubulins are marked on top with the corresponding colors in (c). Large blue (pink) spheres mark the locations in *α*-tubulin (*β*-tubulin) where Kin1 binding induces a large increases in the fluctuations in the residues at the minus (plus) end compared to the fluctuations of the same residues at the plus (minus) end in (Short)_1_-DU-Near (residues with values in the black line above or below the dashed lines). Small pink (blue) spheres denote the locations in *α*-tubulin (*β*-tubulin) where Kin1 binding induces the opposite trends with the large spheres in fluctuations of the residues at the plus and minus ends in (Short)_1_-DU-Near (black line). The response is less asymmetric (value closer to zero) for these small spheres. These residues are visualized respectively in tubulin 3D structure in (c). (c) Side view and top view of the *α*-tubulin (blue) and *β*-tubulin (pink) crystal structures, with neighboring tubulin units (dark grey). Residues with the most asymmetric response are located at the outer surface near interdimer interfaces (large spheres). Residues with less asymmetric response are clustered at the luminal surface and intradimer interface (small spheres). (d-e) Similar to (b) except the difference in the response upon Kin1 binding *κ* is calculated for long protofilament constructs.

The black line in Figure 1(b) shows the difference in tubulin response as a function of the *α*- and *β*-tubulin residues. The data shows that (i) the asymmetric response is transmitted throughout the tubulin structure, and (ii) *α*- and *β* domains within a single tubulin respond very differently. The residues that display the largest asymmetry are highlighted with big blue and red beads in Figures 1(c). Interestingly, these residues are clustered on the outside surface, next to the inter dimer interfaces between the neighboring tubulin dimers. In contrast, residues that display less asymmetric responses (marked with smaller pink and cyan beads in Figure 1(c)) are on the luminal side and the intra dimer interface.

Next, we analyzed the changes in the MT dynamics induced by a Kin1 monomer, which is of particular interest because for processive motion, one of the heads has to be bound to the MT at all times [1, 2, 25, 26, 27, 28, 29]. Thus, to study the effect of a dimeric motor that is in the middle of a step, we bound a single Kin1 head in the middle of a MT segment containing 3 binding sites. Depending on the state of the neck linker, NL, we refer to these constructs as (Short)_1_-U-Near or (Short)_1_-D-Near (Figure 1a). We repeated our calculations to obtain Δ^*R*^ (Equation 9). Interestingly, just as found in (Short)_1_-DU-Near, a single Kin1 motor domain also triggers asymmetric changes in the MT fluctuations with non-zero Δ^*R*^ values (orange and green lines in Figure 1(b)). These calculations show that the plus and minus tubulin dimers respond asymmetrically to the presence of either a single Kin1 head or a Kin1 dimer in the middle binding site(s).

We then constructed a complex with 3 protofilaments (Short)_3_-U-Near to investigate the effect of a bound Kin1 on different potential landing sites for the diffusing head (Figure S3). The response profile of the binding site that would correspond to a side-step is clearly different and almost opposite to the response of the expected binding site.

### Asymmetry in *α*- and *β*-tubulin dimer binding residues

To further analyze the dynamics of the MT residues that directly bind to Kin1 (listed in Table S2 and highlighted in Figure 2(a)), we plot in Figure 2(b) their motilities with and without a bound motor domain for the (Short)_1_-D-Near construct. A comparison of the responses shows a clear differences between the plus- and minus-end binding residues. For example, potential Kin1 binding sites on *β*-tubulin are more mobile at the plus-end but are less so at the minus-end. Conversely, the binding sites on *α*-tubulin are less mobile at the plus-end but are more so at the minus-end. Kin1 binding enhances these differences, showing clear asymmetric effects between the two ends. Since the residues involved in Kin1 binding are mainly positively charged while the residues at the outer surface of the MT are predominantly negatively charged (15 negatively charged residues are shown in Figure 2(a)) [30, 31], it has been suggested that Kin1 diffuses along MT and docks to a particular binding site driven by favorable electrostatic interactions between *α*4 and the negative charges located in the *β* tubulin [32]. We observe enhanced flexibility at the plus-end of the main Kin1 binding domain (H12 of *β*-tubulin), which could lead to a stronger binding affinity, implying that the plus-end should have a stronger interaction with the diffusing head than the minus-end.

**Figure 2:**
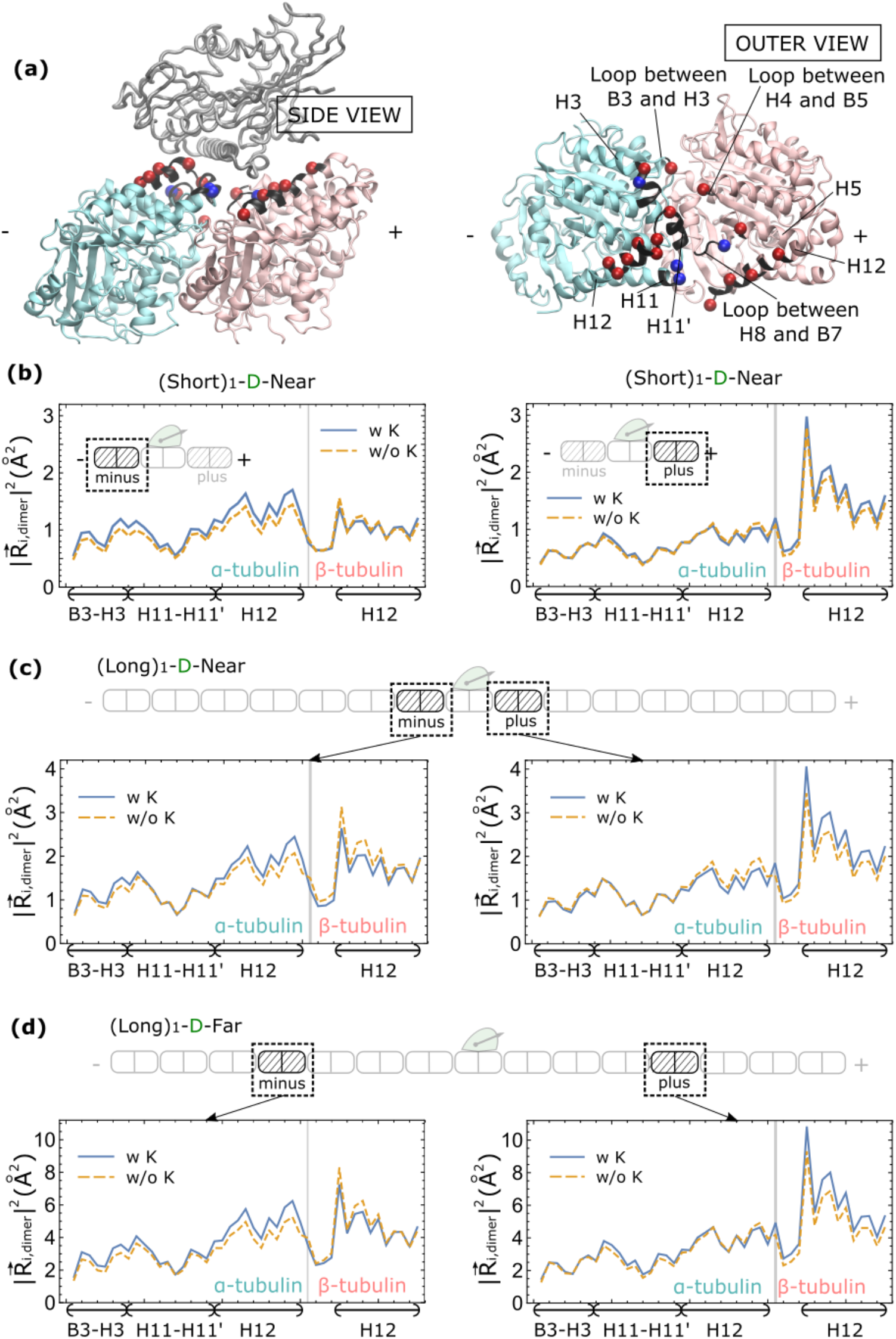
The asymmetric response of Kin1 binding sites of plus- and minus-end tubulin. (a) Side view (left) and outer view (right) of the *α*-tubulin (light cyan) and *β*-tubulin (pink) structures with a bound Kin1 trailing head (gray). Potential Kin1 binding sites (listed in Table S2) are highlighted in black, their charges are shown in red (negative, 15 total) and blue (positive, 4 total). (b-d) Mobility relative to the center of mass of the corresponding dimer 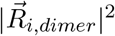 (Equation 5) of potential binding sites at the minus-end (left) and plus-end (right) of the tubulin with (dotted line) and without (solid line) the bound Kin1 monomer in the (Short)_1_-D-Near (b), (Long)_1_-D-Near (c) and (Long)_1_-D-Far (d) structures, respectively.

### Stresses generated by Kin1 binding expands the MT lattice

We next investigated the effect on the elasticity of the MT lattice when Kin1 binds by comparing 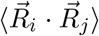 and 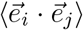 of tubulin dimers with (upper triangle matrix) or without Kin1 (lower triangle matrix) in Figure 3(a-b). The sequence of each tubulin is divided into three domains, highlighted in the 3D structures in Figure 3(c-d).

**Figure 3:**
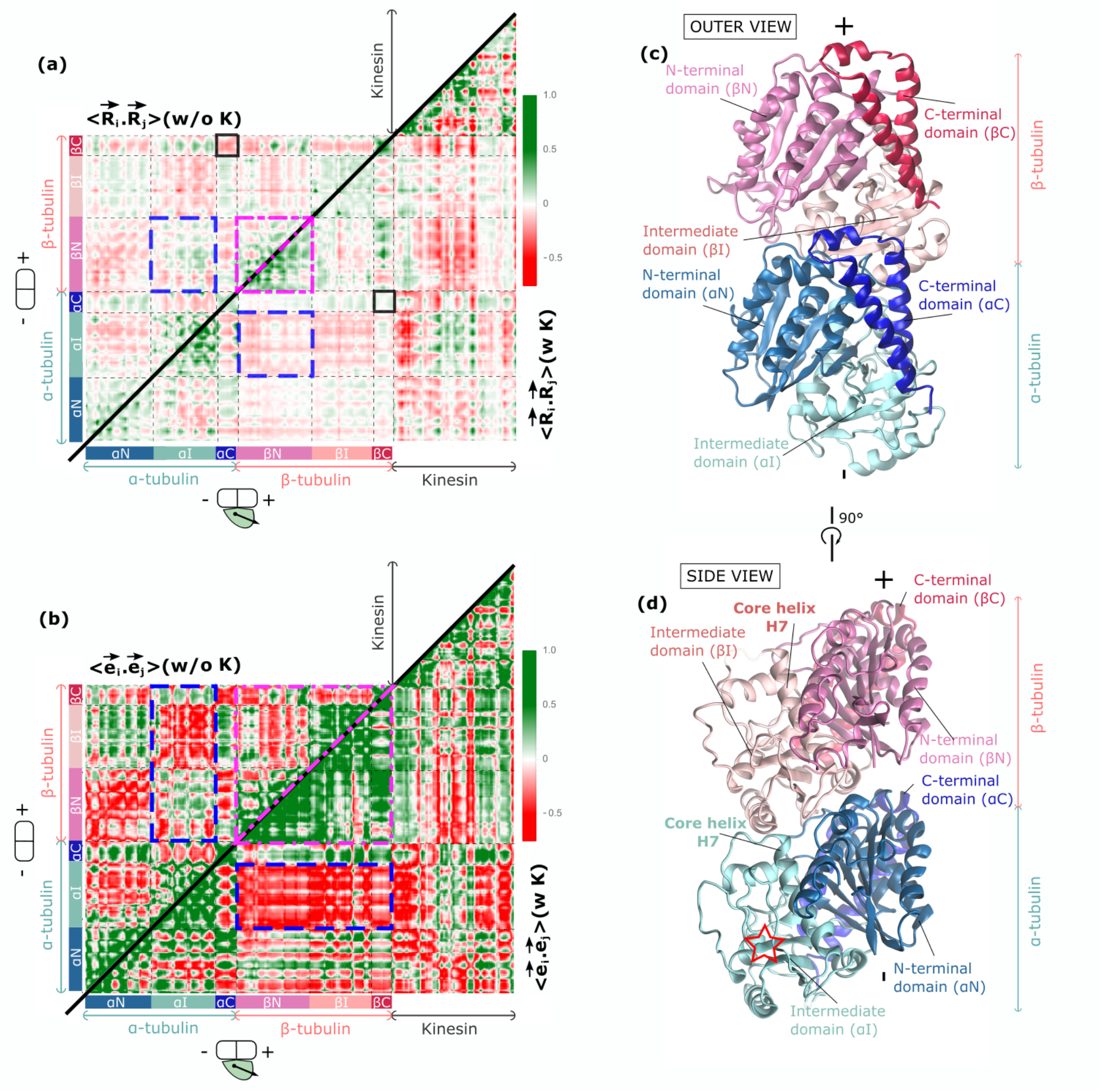
Correlations between residues in tubulin before and after Kin1 binding. (a) Plots of spatial 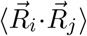 and (b) orientational 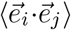 correlation matrices for an *αβ*-tubulin dimer without a Kin1 (upper matrix, labelled top left corners) and with the bound trailing head (lower matrix, labelled side right corners). The plots are on the same color scale with green (red) representing positive (negative) correlation. Note that any value equal to or greater than 1 is shown in dark green. Each tubulin has 3 domains [35]: an N-terminal domain (*α*N or *β*N), an Intermediate domain (*α*I or *β*I) and a C-terminal domain (*α*C or *β*C). The corresponding residues are marked as bars with different colors. Boxes with different colors highlight regions with maximum changes between the residues with and without bound Kin1. (c) Outer view and (d) side view of the *α*- and *β*-tubulin structure with different tubulin domains, colored corresponding to (a) and (b). A red star in (d) highlights the position of the *α*I domain, which is in the same position with the most increase in mass, detected in Cryo-EM data upon Kin1 binding to the MT (Figure 6 of [13]).

The correlation matrix, 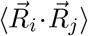, shows that Kin1 induces coordinated motion involving key residues within the tubulin dimers. First, two small black squares in Figure 3(a) represent the effect of switching from anti-correlated motion, in the absence of Kin1, to positive correlation when Kin1 binds to MT between the C-terminal domains of the *α*- and *β*-tubulins (*α*C and *β*C) (see also Figure 4(b)). Second, the enhancement in the correlations is greater within the N-terminal domain of the *β*-tubulin (*β*N vs. *β*N, see the two connected magenta dash-dot triangles, the triangle below is greener). Third and most interestingly, the intermediate domain of the *α*-tubulin, without direct contacts with Kin1, becomes negatively correlated with the N-terminal domain of the *β*-tubulin (*α*I vs *β*N, see the two blue dashed rectangles).

**Figure 4:**
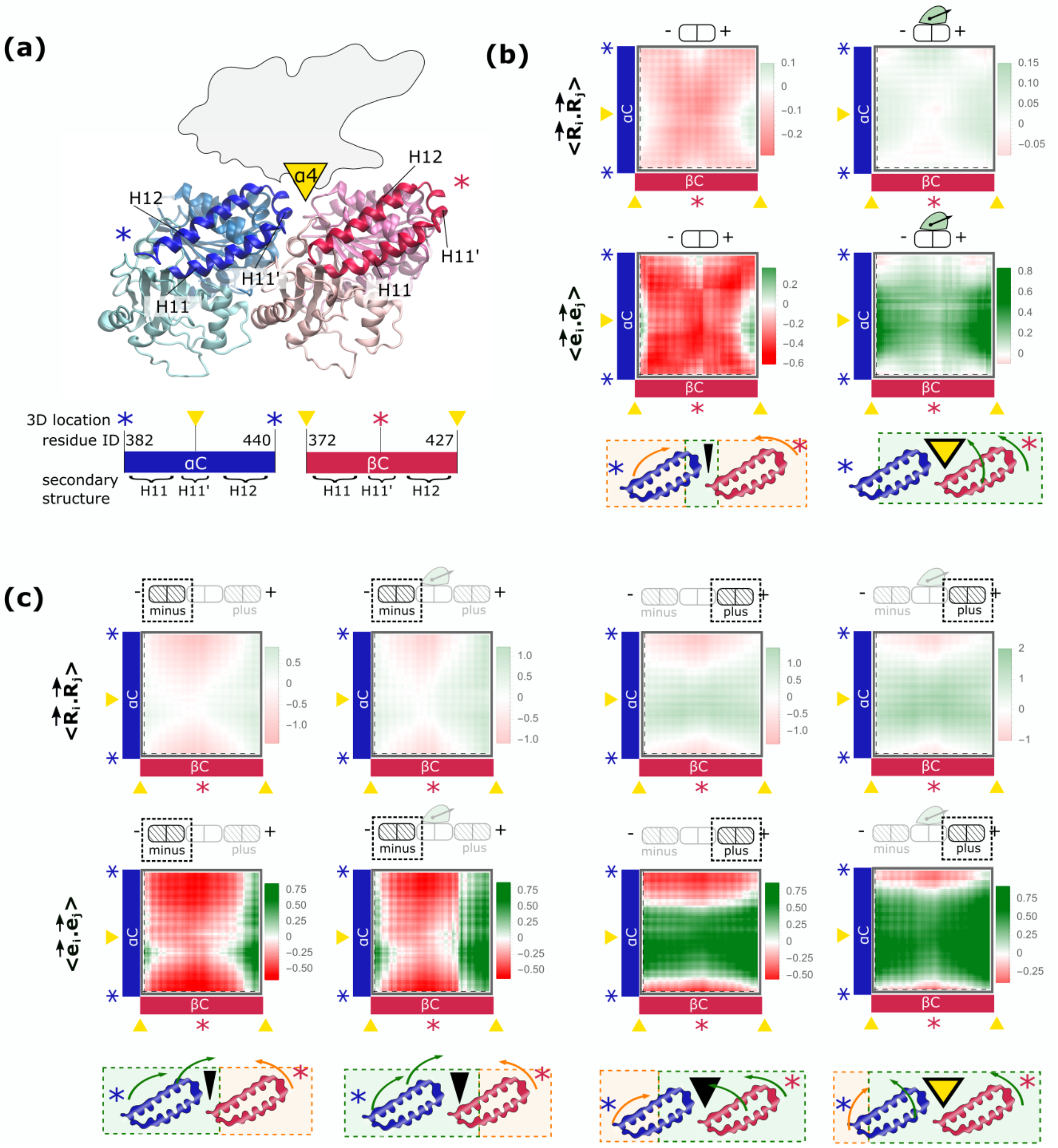
The response of the outer surface of MT to Kin1 binding. (a) Top: Side view of a tubulin dimer with C-terminal domains of *α* and *β* tubulins (*α*C and *β*C, highlighted with darkest blue and red colors), consisting of H11, H11’ and H12 helices [35]. Upon binding, helix *α*4 in Kin1 (shown as a yellow triangle) is between the two C-terminal domains and at the intradimer interface cleft. Bottom: sequences of *α*C and *β*C are represented as blue and red bars, respectively. The locations of the residues near the intradimer and interdimer interfaces are marked with yellow triangles and stars, corresponding to the top figure. (b) Plots of 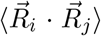 (upper panels) and 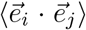 (middle panels) between C-terminal domains of an *αβ*-tubulin dimer in the absence of Kin1 (left panels) and with the bound trailing head (right panels). The plots are on the same color scale with green (red) representing positive (negative) values. Bottom panels: The positions of the residues with positive correlations are mapped to side views of corresponding C-terminal domains with a dashed green boxes; the negative correlations are in orange boxes. (c) Similar to (b) but correlation matrices are calculated for minus- and plus-end tubulin dimers in the (Short)_1_-D-Near construct.

Remarkably, the very act of Kin1 binding results in substantial conformational changes in the tubulin dimers, which results in the distortion of the MT lattice. In particular, Kin1 binding leads to a more stable *β*-tubulin and twisted *α*-tubulin, which is reflected in 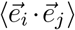 in Figure 3(b). In the absence of Kin1, there are no apparent alignments in the dimers, represented by patchy colors throughout the upper matrix. Upon Kin1 binding (lower matrix), the entire *β*-tubulin is aligned with each other (most values are green within the highlighted magenta dash-dot triangle in the lower matrix). The soft modes show that the motion is anti-correlated with the intermediate domain of the *α*-tubulin (most of the values are red in the highlighted blue dashed rectangles in the lower matrix).

These results are consistent with the previous observation that although *β*-tubulin is the primary binding site of Kin1, most differences between the Cryo-EM data of MT coated with and without Kin1 are found in the *α*I domain of *α* tubulin [13]. Kin1 binding results in an increase in the mass near the center of the *α*-tubulin towards the binding interface of the adjacent tubulin-dimer in [13]. This increase in mass (marked with a red arrow in their Figure 6A, 6F and 6G of reference [13]) is at the same location of the *α*I domain (marked with a red star in our Figure 3(d)). From our calculations, the most striking effect of Kin1 is the anti-correlated fluctuations between the entire *β*-tubulin and the *α*I domain, which increases the mass due to co-localization in this region. Recent Cryo-EM data on different types of MTs also show that changes in the MT lattice arise mainly due to the changes in *α*-tubulin [33, 34]. The negative correlation between the intermediate domain of *α*-tubulin and the *β*-tubulin means that the two parts undergo collective movement that oppose one another, which could distort the polar track. As a consequence, the MT lattice expands upon Kin1 binding [15, 14].

### Kin1 dramatically alters the soft modes of the C-terminal domains of tubulin

We then focus on helices H11, H11’ and H12, classified as the “C-terminal domains” [35] (see also Figure 4a). The C-terminal domains include several Kin1 binding sites with many that we identified earlier as having the largest values of Δ^*R*^ (see Figure 1c). Figure 4b shows two distinct correlation profiles, 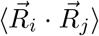 (dynamic) and 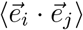 (orientation, see Methods). The C-terminal domains of the two tubulins are *anti-correlated* (red) when Kin1 is not bound, they are *correlated* (green) when a Kin1 is bound. The two distinct profiles show that, upon binding, residues in the MT binding regions of kinesin become stiffer (see Figure S1 in the supporting information) with *α*4 helix of Kin1 wedging into the cleft formed at the intradimer interface between the two C-terminal domains, keeping the cleft open (represented by a yellow triangle in (Figure 4(a)). On the other hand, when Kin1 is not bound, the cleft is empty, allowing anti-correlated soft modes to be dominant in the outer surface of tubulin, effectively closing the cleft. The two-state closed to open transition expands the MT lattice, and creates the asymmetric response that sets the stepping directionality.

### Kin1 opens a cleft at the *α*/*β* intradimer interface and enhances binding at the MT plus-side

Having determined the distinct correlation profiles of the Kin1-bound and Kin1-free modes at the binding site itself, we tested the correlations at the adjacent binding sites as well. We expect that correlated motions between the C-terminal helices would increase the probability that additional Kin1s would bind to the plus end tubulin since they keep the cleft open (referred to as open cleft (**OC**)) while anti-correlated movement that effectively closes the cleft (**CC**) would decrease the binding probability. We calculated the correlation profiles of the plus and minus ends binding sites when Kin1 is absent as well as when it is bound to the middle binding site of the (Short)_1_-D-Near structure. The right and left sides of Figure 4(c) profiles are dramatically different at the plus and minus ends of the MT. In particular, upon Kin1 binding, the correlation profiles of the plus-end binding site (the right-most column of Figure 4(c)) are similar to the profiles of the tubulin-Kin1 complex in Figure 4(b). Although Kin1 binding extends the correlated region in the C-terminal domains of the minus-end tubulin dimer, their motions remain mostly anti-correlated, suggesting that the cleft between the two dimers is effectively closed. The relative correlations between the residues adjacent to the cleft determine Kin1 binding affinity. In the region of a MT that is on the plus-side of an already bound Kin1 cleft is correlated and effectively open with a high affinity for Kin1 binding. Thus, the probability of the cleft being open to the right (plus end of the MT) of bound Kin1 is higher than in the minus end, which is a cooperative effect that propagates (see below).

### Asymmetric plus and minus response is a long range effect

To investigate whether the propensity for the cleft to in the **OC** state, poising another Kin1 to bind, can be sustained over long distances, we tested the effect of a bound Kin1 monomer and dimer on the MT dynamics by incorporating larger MT segments in constructs shown in Figure 1(d-e) and Figure 2(c-d). In all cases, the dynamic responses of the MT were alike. Surprisingly, there is an enhanced probability for the tubulin dimers in the plus end of a bound Kin1 to be in the **OC** state compared to the minus end. The cooperative transition from Kin1 induced **CC** → **OC** persists even several tubulin dimers to the plus end of the location of the bound Kin1 (Figure 1(e)). Because of the two-state closed to open transition, the long-range strain-induced asymmetric response of the MT to Kin1 binding could occur in a one dimensional lattice even if the interactions are short-ranged.

### Microtubule binding induces correlations within the nucleotide binding sites in the motor

So far we have only considered the changes in the MT dynamics and structures in the *α/β* tubulin upon Kin1 binding. It is appreciated that the MT has a profound effect on the steps in the catalytic cycle of the motor. For instance, MT enhances ADP release rate by nearly a factor of 1,000 [12]. In addition, it has been proposed that cooperative motion of switches I and II is necessary for many steps in the biochemical cycle [36, 37]. We tested the possibility that binding to MT induces correlated response in the relative dynamics of switches I and II, along with ADP and its binding sites, which might provide a partial explanation for the significant ATPase rate changes.

In order to assess the changes in the fluctuations of switches I and II, we analyzed the correlations between the switches for a kinesin bound to a MT, and for the unbound motor. We used the 30 lowest energy normal modes of an ADP-bound monomeric kif1a (1i5s.pdb [38]), and an ADP-bound kif1a-MT complex (2hxh.pdb [39]). For both the structures, we calculated the average dynamical correlations, 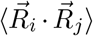, between all the pairs of residues in switches I and II (Equation 6 in the Methods section). Figure 5(a) compares the two correlation matrices for MT-bound-kinesin (bottom) versus the unbound-motor (top – plotted with the same color scale with the bottom panel). A greater value with darker green indicates an increase in correlation between a given pair of residues in the switches. There is a remarkable difference in 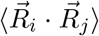 with and without MT. There is a significant increase in the correlations upon MT binding for all the residue pairs, with almost the entire matrix becoming nearly dark green (compare the bottom panel to the top in Figure 5(a)). For example, binding to a MT enhances the correlations between the fluctuations of kinesin residues R216 and E253 (highlighted in the black boxes) by a factor of ≈ 5.

**Figure 5:**
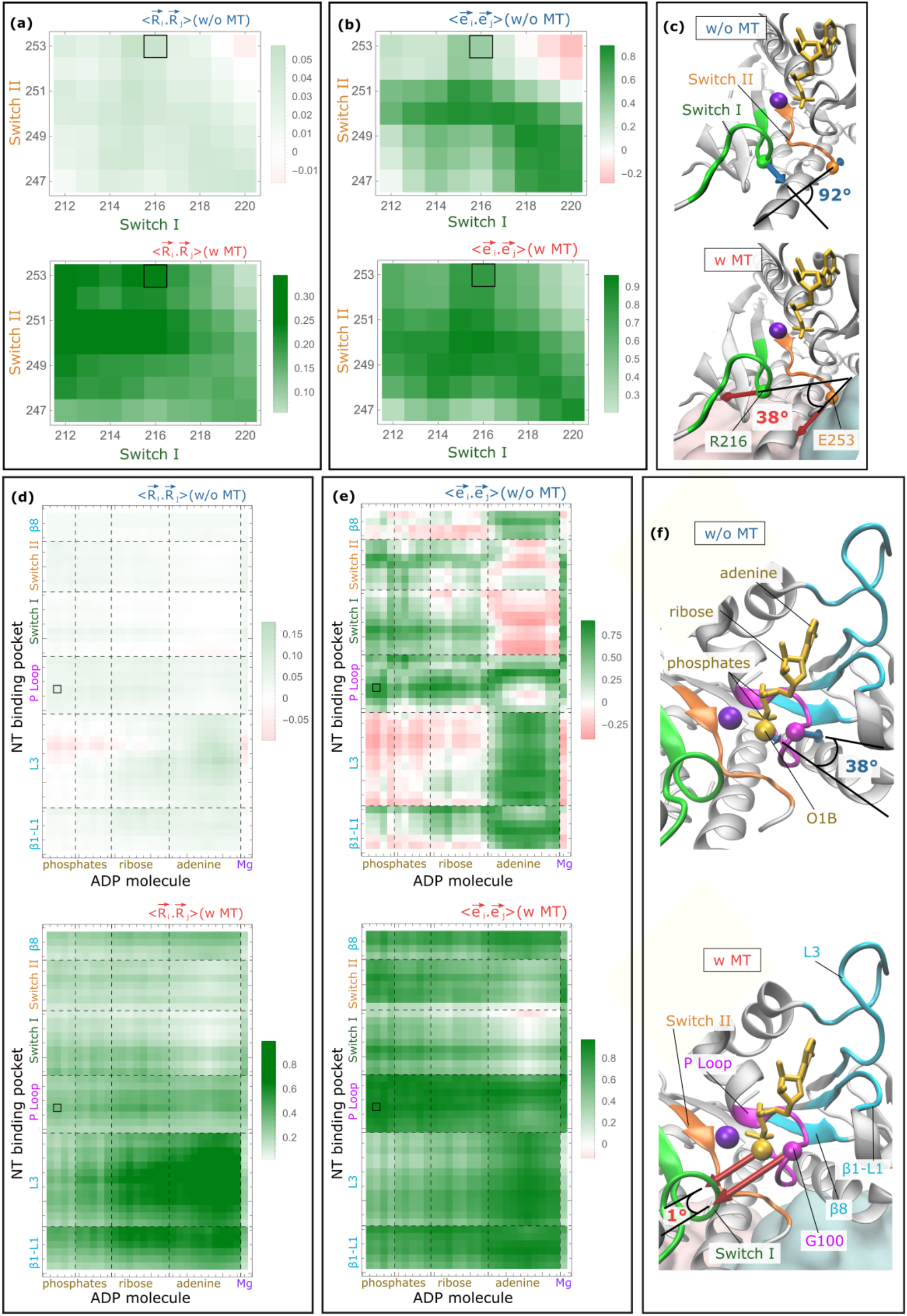
(a-c) MT binding induces correlated translational and directional movements between switch I and switch II. (a) Spatial correlation matrices 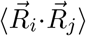 and (b) orientational correlation matrices 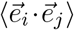 (labelled on the top right corner) between switches I (residues 212-220) and II (residues 247-253) for the unbound kinesin 1i5s.pdb (top panels) and the MT-bound-kinesin 2hxh.pdb (bottom panels). Positive correlation is shown in green hue and negative correlation is shown in red hue (scale bars on the right). (c) Structures of switch I (green) and switch II (orange) of ADP binding site relative to ADP molecule (yellow) and Mg^2+^ ion (purple) when the kinesin motor is free (top), or bound relatively to an *αβ* tubulin dimer (light cyan and pink) (bottom). Residues R216 (switch I) and E253 (switch II), marked with stars in the 1i5s sequence in Figure S1(d), are represented as green and orange beads at the *C_α_* atoms. Blue (top) and red (bottom) arrows are the first eigenvectors, corresponding to the lowest energy mode. (d-f) Correlation between ADP and the binding site residues increases upon MT binding. (d) Spatial correlation matrices 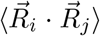 and (e) orientational correlation matrices 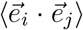 between ADP and the binding residues when kinesin is free (top) and bound to MT (bottom). (f) ADP (yellow), Mg^2+^ ion (purple) and the ADP binding site (color-coded accordingly as in (d-e)) in the unbound kinesin (top) and MT bound kinesin (bottom). Residue G100 of P loop, marked with a magenta star in the 1i5s sequence in Figure S1(d), and O1B atom of ADP are represented as yellow and magenta beads at their alpha carbon atoms, respectively. Their motion is represented using the first eigenvector in the lowest energy mode shown (blue and red arrows). The angle measures the relative motion between the two elements.

There are two independent factors that might contribute to the enhancement in the correlated motions reflected in 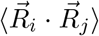 in the presence of the MT: (i) the increase in the spatial fluctuations, and (ii) directional alignment in the residue motions. In order to isolate these effects, we calculated the average orientational correlations 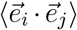 between all pairs of switch I and switch II residues (Equation 7). Figures 5(b) (plotted with the same color scale) show the directional correlation matrices in the two cases. Almost all the values in the matrix plotted for MT-bound-kinesin are larger (darker green color in Figure 5(b) bottom) than for apo-kinesin (Figure 5(b) top), indicating that the fluctuations in the two switches are orientationally aligned upon binding to MT. The correlations between residues R216 and E253 (black boxes) increases by a factor of two upon MT binding. Comparisons of the results in the two panels in Figures 5(b) shows that the extent of alignment of the switches I and II, as measured by 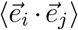 increases when the motor binds to the MT. In Figure 5(b), the bottom panel shows that the residues in the two switches exhibit a high degree of orientational correlation with complete absence of anti-correlation, observed in the upper right corner in the top panel.

To visualize the changes in the movements of the two switches, we plot in Figure 5(c) the lowest energy eigenvectors associated with residues R216 and E253 of the isolated kinesin and a MT-bound kinesin. For an unbound motor (the top panel) the eigenvectors of the pair are almost perpendicular. In contrast, upon binding to a MT (the bottom panel) the same residues are significantly aligned with each other – the relative angle of the first eigenvectors changes from 92° to 38° accompanied by increased fluctuations. Taken together, these results show that interactions with the MT produces correlated responses in key structural elements in the motor domain.

To obtain additional insights into the increase in the ADP release, we analyzed the correlations using 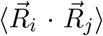 between the fluctuations in the nucleotide and the residues in the binding pocket. Comparing the correlations calculated using a MT-bound and an unbound kinesin in Figure 5(d), we find that most of the values in the bottom panel of Figure 5(d) are greater (represented in dark green) than in the top panel, indicating an increase in the dynamical correlations upon MT binding.

The increase in 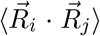 upon binding to MT is caused by an increase in the magnitude of the fluctuations, as well as an increase in the orientational correlations 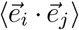. The results in Figure 5(e), showing the relative changes in the directional correlations upon MT binding, establishes that binding to the MT induces alignment in the fluctuations of the nucleotide binding pocket and the ADP molecule. Without MT, different structural components of the ADP molecule show opposite fluctuations with parts of the binding pocket that they do not directly interact with each other. For instance, adenine does not directly interact with the switches (see Figure 5(f)), and they have negative 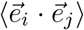 values (represented in red hue in Figure 5(e) top panel). Similarly, Mg^2+^ ion, ribose and phosphate are far from from L3 (see Figure 5(f)), and they also have negative 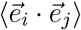 values (red hue in Figure 5(e) top panel). However, upon MT binding, ADP is aligned with the whole nucleotide binding site, as shown by the green hue in Figure 5(e) bottom panel. Hence, MT binding results in the entire nucleotide binding sites to fluctuate in unison, which could explain the faster release of ADP.

As switches I and II harbor the Mg^2+^ ion and the *γ*-phosphate, another conserved motif, normally referred to as the P loop, stabilizes the *α*- and *β*-phosphates of the nucleotide molecule. To illustrate the ADP fluctuations, we considered the correlations between residue G100 in the P loop and ADP phosphate atom O1B. We highlighted their values with black boxes in Figure 5(d-e), and illustrate their first non-zero eigenvectors in Figures 5(f). After binding to MT the amplitudes of fluctuations increase more than 3-fold in both G100 and O1B. In addition, there is enhanced directional alignment (Figure 5(f) bottom versus top). We surmise that MT binding significantly alters the local dynamics of the nucleotide binding pocket by making the constituent residues fluctuate with larger amplitude along the same direction.

We also calculated the 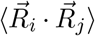 and 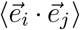 between AMPPNP and the binding pocket before and after MT binding (data not shown). We found that, just as is the case for ADP binding, both 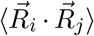 and 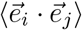 between AMPPNP and the binding pocket increase upon MT binding. It is possible that the increase in the correlation between the fluctuations of ATP and its binding pocket contributes to an increase of ATP hydrolysis rate upon MT binding [40].

These findings are consistent with two experimental observations. First, a recent structural study [41] shows that Mg^2+^ inhibits ADP release in kif1a and that the fluctuations (measured by B-factors) of ADP increase during Mg^2+^ release. The increase in the fluctuations of ADP upon MT binding, found here, may similarly correlate with the rate increase in ADP release enhancement. Second, EPR spectroscopy of spin-labelled ADP [42] showed that the relative angular motions (as measured by a cone angle) between ADP and its binding pocket decrease upon MT binding. This is in accord with our finding that the directional correlation, 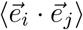, between ADP and the associated binding pocket increases upon MT binding, which is manifested as a decrease in the cone angle.

### Reciprocal interactions between kinesin and MT, processivity and ADP release

The asymmetric response of the tubulin at the plus and minus ends of the MT, expansion of the mechanical stress induced MT lattice, and the **CC** → **OC** transition leads to a conceptual model that we illustrate in Figure 6. The expansion of the MT lattice, accompanied by the formation of the **OC** state in which the fluctuations in the *β* domain are enhanced, propagates preferentially towards the plus end of the MT. The consecutive formation of the **OC** states to the right (towards the plus end of the MT) of a bound Kin1 (schematically indicated by the green region in Figure 6) leads to preferential binding of additional Kin1s towards the plus end. The same picture holds when only a single motor, which are most commonly studied in *in vitro* single molecule experiments, walks processively on the MT. In this case, when Kin1 binds it induces multiple **CC** → **OC** transitions in the tubulin dimers that are at the plus end. When the trailing head detaches and diffusively searches for the subsequent binding site, it is more poised to bind to the **OC** than to the **CC** state, which would ensure processivity. Thus, it is the mechanical stress-induced formation of the **OC** states, which although occurs within a single *α/β*-tubulin but has a long range effect, that explains both the directionality and processivity of kinesin motors. Conversely, *α/β*-tubulin alters the motor dynamics, especially at the nucleotide binding sites (Figure 5), which likely facilitates the release of ADP.

**Figure 6:**
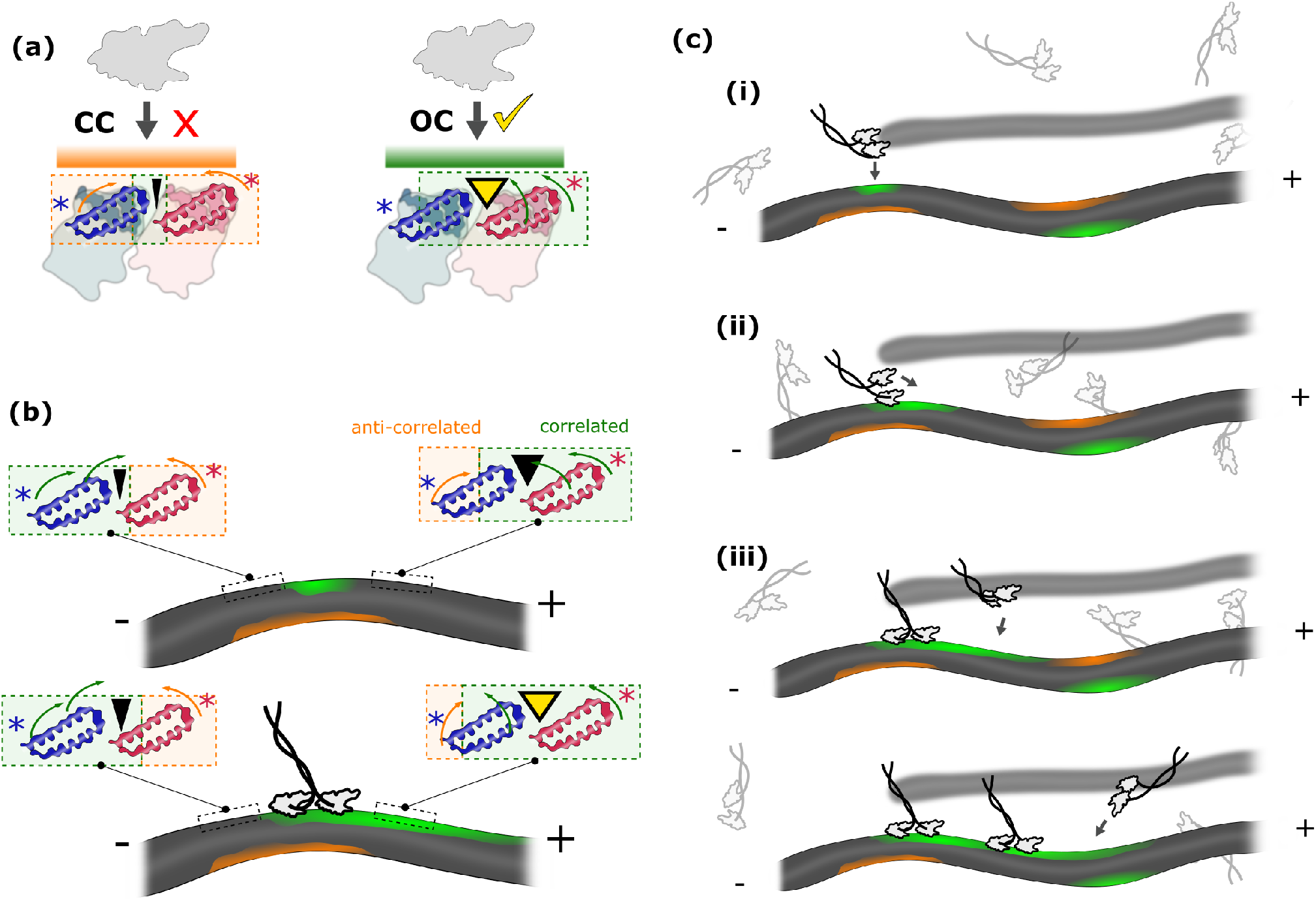
(a) Two soft modes characterize the dynamics of the (**CC**) and stress-induced (**OC**) states at the cleft of the *α/β* C-terminal domains: (left) anti-correlated movements between the two C-terminal domains which effectively close the cleft, inhibiting Kin1 binding (marked by the red cross); (right) correlated movements across the entire Kin1 binding site which enable the docking of *α*4 helix into the open cleft at the intradimer *α* − *β* interface. The more and less Kin1 binding competent outer surfaces, corresponding to the **CC** and **OC** states, are marked with orange and green, respectively, in Figures (b-c). (b) Kin1 binding induces correlated soft modes in the tubulin dimers at the plus end, resulting in the formation of **OC** states in the plus-end direction for other motor head(s) to bind with high probability. (c) Schematics of effects of the asymmetric response inside cells. (i) Initially Kin1 binds with high probability to MT sites with open clefts (marked in green). The green regions form at the convex side of a bent MT [15]. (ii) As one of the motor binds to the MT, it changes the soft modes of the surrounding binding sites, making the adjacent binding site at the plus end of the protofilament more suitable than at the minus end for the binding of the diffusing head. This has the effect of minimizing the probability of the motor taking side or back steps. (iii) A bound Kin1 dimer induces soft modes in tubulin dimers at the MT plus end to be more correlated than at the minus end, thus attracting additional motors to bind to the plus end of the same MT. This creates a cooperative effect that promotes the other motors to land on a specific protofilament MT while avoiding binding to other protofilaments (represented with a blur MT in the background) at non-saturated Kin1 concentrations.

## Conclusions

In this paper, we specifically focus on the interactions between kinesin and MT, how MT dynamics impacts the stepping of the motor, and how, in return, the motor impacts the internal dynamics of MT. Here, we pieced together related tubulin and different MT bound Kin1 structures, along with MT bound kif1a and free kif1a structures with ADP to create models in various states. The residue-dependent mechanical responses show that the interface between the MT and the motor stiffens upon binding, which in turn allosterically alters the proximal nucleotide binding sites. The MT induces correlations between switch I and switch II, and ADP by aligning the binding sites along a preferred orientation. The internal dynamics of the motor as well as the MT track reveal that interaction between Kin1 and the MT produces a long-range (possibly across the whole length of the MT) allosteric effects that govern the reciprocal dynamical relationship between MT and the motor.

The most important finding of this work is that the plus and minus ends of the MT respond asymmetrically to Kin1 binding to a specific *α/β* tubulin dimer. Surprisingly, the asymmetry persists over a long spatial range, which is made possible because Kin1 induces stress-induced closed (**CC**) to open (**OC**) transition in the cleft of the *α/β* C-terminal domains, the binding site of the motor. The **CC**→ **OC** transition also expands the MT lattice [15, 14], presumably downstream of the site of the bound motor. The accessible kinesin binding sites in the front and rear ends of *α/β* tubulin produce asymmetric mobility patterns (Figure 2). The dominant soft modes of the MT-Kin1 complex results in the binding site being correlated, which persists on the plus-side of the MT but not on the minus-side (Figure 4 and 6a-b). Thus, the inherent bias in the stepping towards the plus end of the MT is encoded in the strain-induced change in the affinity of the motor for sites on the *α/β* dimer. This principal finding, whose origin is mechanical in nature, rationalizes the experimental observation [24] in structural terms. Most importantly, it implies that binding of an individual motor induces long-range correlations in the MT lattice [24, 43], which leads to preferential binding of kinesin at sites to the plus end of an already bound motor.

Our calculations also show that Kin1 induces positive correlations within the *β*-tubulin, its primary binding monomer, and anti-correlations between the *β*-tubulin and the intermediate domain of *α*-tubulin – the *α*I domain (Figure 3). This is in excellent agreement with the CryoEM [13]. The recent CryoEM data [34, 44] observed that MT lattice can be in 2 different states, with the GDP-MT lattice is more compact than the GMPCPP-MT lattice. The expansion/compaction arises from the rearrangement of the *α*I domain, which is in accord with our calculations. Furthermore, it has been shown Kin1 can expands the compact GDP-MT lattice [15, 14], suggesting that the Kin1-bound GDP-MT structure should have a higher pitch distance closer to the GMPCPP-MT lattice state in solutions.

We interpret the origin of the **CC** → **OC** transition, which is linked to the MT expansion, to the dramatic alterations in the soft modes within the tubulin dimers. This affects the Intermediate domain *α*I to move in an anti-correlated manner relative to the entire *β*-tubulin. These highly anti-correlated modes redistribute the mass inside *α*-tubulins [13], thus effectively expanding the GDP-MT lattice *in solution* [15, 14]. The extensive dynamical interplay between the motor and MT, shows that MT should be thought of as an active, and dynamic material. Indeed, the most important conclusion from our study is that it is the interplay of the conformational dynamics involving both kinesin and the tublin dimers that sets the stepping direction. This results in anisotropic response in tubulin when kinesin binds that could propagate in an asymmetric manner throughout the MT lattice.

## Methods

### Elastic Network Model (ENM)

We model kinesin and MT as networks composed of N nodes with each residue representing a single node, located at the position of its C_*α*_ atom. In the calculations with explicit ADP (AMPPNP), each nucleotide heavy atom is also taken to be a node. Two nodes, *i* and *j*, are connected by a harmonic spring if the distance between them in the Protein Data Bank (PDB) structure is less than *R_c_* =10 Å. We have verified that changing the cut-off distance over a reasonable range up to *R_c_* =8 Å does not qualitatively alter the results.

The potential energy function for the elastic network is [20, 21, 22],

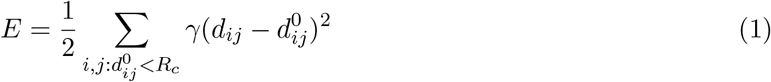

where *γ* is the spring constant that defines the energy scale (we set to be one unit), *d_ij_* is the instantaneous distance between residues *i* and *j*, and 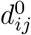 is the corresponding PDB distance, which is assumed to be the equilibrium separation. The dynamics of the system around the equilibrium state (the PDB structure) is determined from the normal modes, which are calculated by diagonalizing the Hessian matrix **H** obtained from the second derivatives of the internal energy in Equation 1. The eigenvalues of the matrix, *λ_m_*, correspond to the energies of the system and the fluctuations of the modes are calculated from the corresponding eigenvectors *q_m_*. The eigenvectors can be expressed in terms of the Cartesian components of the fluctuations of individual residues as,

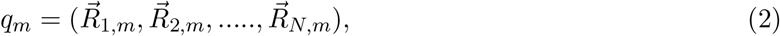

where 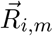 are the (*x, y, z*)-components of the *i^th^* residue in motion. In practice, we used 30 lowest energy normal modes in our calculations. Our results remained unchanged when we increased the number of modes (see SI).

### Local stiffness, correlations between residues, and MT response

In order to determine the local stiffness, *κ_i_*, we used the Structural Perturbation Method (SPM) [18, 19]. The SPM, is used to probe the response of all the residues in response to a local perturbation at a specific residue. Within the ENM (Equation 1) the local perturbation is realized by changing the value of *γ*. The perturbation alters the elastic energy in the normal modes. We define the local stiffness, *κ_i_*, as the fraction of the effective elastic energy (SPM) stored in the *i^th^* residue over the residue fluctuation amplitude squared (the mobility), which turns out to be,

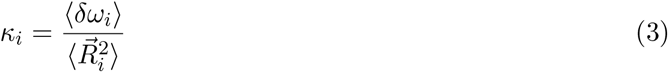

where the mobility 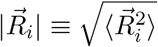 of a residue is,

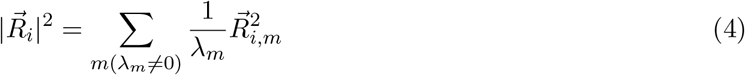

and 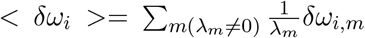 with *δω_i,m_* is the SPM response of residue *i* at mode *m*. The residue-dependent alterations in the stiffness and their locations in the structures are illustrated in Figure S1.

To separate the fluctuations of residues in a dimer from the dominant global movement, we determined the global mode of dimer *d* as 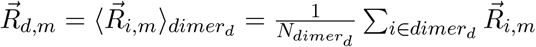 with 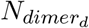 is the number of residues in dimer *d*. The motility of a given residue relative to the center of mass of the corresponding dimer is:

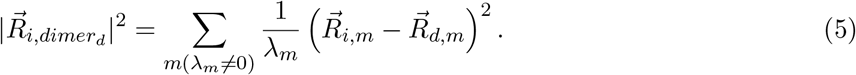

We define the average dynamic correlation between residues *i* and *j* as,

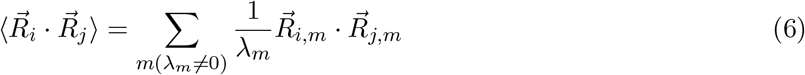

where 〈…〉 implies an average, ideally taken over all the modes. In this paper, we use the first 30 lowest modes, which are the most dominant. In addition, we calculated the directional correlation between residues *i* and *j* using,

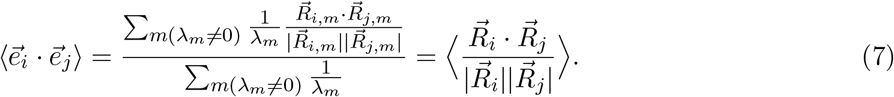

The dynamic response, 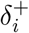, of a plus-end MT residue to kinesin binding is defined as,

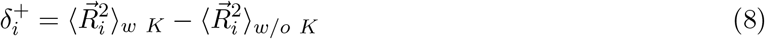

where *w K* (*w/o K*) is calculated with (without) the bound kinesin, and the superscript refers to the plus end of the MT. The minus-end response, 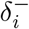, can be similarly defined. The asymmetric nature of the response can be assessed using

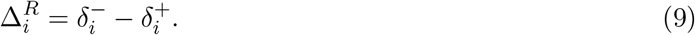

If the response of MT to kinesin binding is symmetric then Δ^*R*^ in Equation 9 would be zero, which implies that a non-zero value would indicate an inherent asymmetric response.

### Kin1 dimer and MT structure preparation

The Kin1-MT protofilament structure preparation consisted of several steps, which are similar to the procedure described elsewhere [7]. First, we extracted a protofilament structure from the structure of a segment of the MT [45]. We augmented the filament with additional *αβ*-tubulin (1jff.pdb [46]), segments as needed. Next, we docked the two new kinesin-tubulin structures provided in the PDB (4uxy and 4uxt [47]) to two adjacent binding sites on the MT filament so that the relative positions of the kinesins on the tubulin were similar to the solved kinesin-tubulin complexes. The leading head has a disordered NL and the trailing head has a docked NL. We repeated our calculations with an averaged equilibrium structure of the dimer (with full NLs) bound to the MT obtained in our previous study [7], obtaining similar results, thus establishing the robustness of the calculations. For the construct of single kinesin head bound with docked or undocked NL to *α/β*-tubulin heterodimer, we used either one of the two PDBs (4uxy or 4uxt).

## Acknowledgements

We thank Naoto Hori, Mauro L Mugnai and Robert A Cross for fruitful discussions. DT is grateful to the National Science Foundation (CHE 19-000033) for support. Additional support from the Welch Foundation, through the Collie-Welch Chair (F-0019) is acknowledged. HTV is currently funded through an EMBO Long-term Postdoctoral Fellowship (EMBO ALTF 904-2019).

