## Supplementary Information for "Plus and minus ends of microtubule respond asymmetrically to kinesin binding by a long range directionally driven allosteric mechanism"

Table S1: Residues with the high stiffness in *apo* kinesin (1i5s.pdb) and MT-bound kinesin (2hxh.pdb). The residues are also shown in Figure S1(b) and (c).

| *Apo* **kinesin**<br>**Residue ID in 1i5s.pdb** | **Structural Element** | **MT-bound kinesin**<br>**Residue ID in 2hxh.pdb** |
|---|---|---|
| 56, 59 | $\beta$2-L3 | |
| 97-98, 100 | P loop | 98 |
| 107-108 | $\alpha$2a | |
| 110-111, 115-116 | L5 | 116 |
| 120 | $\alpha$2b | |
| 148-150, 155 | $\beta$4 | |
| | $\beta$5a | 170-171 |
| | $\beta$5b | 175 |
| 202 | $\alpha$3 | 202 |
| 213-220 | Switch I (L9 & $\beta$6) | 211, 213-216, 219 |
| 249-254 | Switch II (L11) | 250, 252-255 |
| 268 | $\alpha$4 | 265-269, 271-272, 276-278, 280 |
| | L12 | 304-305, 307-308 |
| 313, 315 | $\alpha$5 | 311, 314-317 |
| 332 | L14 | |
| 341-342, 345, 347 | $\alpha$6 | 339, 342-347 |
| $Mg^{2+}$ | | $Mg^{2+}$ |

Table S2: Potential kinesin binding sites on the MT.

| **Residue ID in 1jff.pdb** | **Structural Element** | **Note** |
|---|---|---|
| 97 | Loop between B3 and H3 of $\alpha$-tubulin | GLU |
| 113 | H3 of $\alpha$-tubulin | GLU |
| 411 | H11' of $\alpha$-tubulin | GLU |
| 414-415, 417, 420, 423 | H12 of $\alpha$-tubulin | GLU |
| 424 | H12 of $\alpha$-tubulin | ASP |
| 163 | Loop between H4 and B5 of $\beta$-tubulin | ASP |
| 194 | H5 of $\beta$-tubulin | GLU |
| 420[a], 431[a] | H12 of $\beta$-tubulin | GLU |
| 427[a], 437 | H12 of $\beta$-tubulin | ASP |
| 112 | H3 of $\alpha$-tubulin | LYS |
| 401-402 | H11 of $\alpha$-tubulin | LYS |
| 262 | Loop between H8 and B7 of $\beta$-tubulin | ARG |
| 108-111, 114 | H3 of $\alpha$-tubulin | |
| 400 | H11 of $\alpha$-tubulin | |
| 405-410 | H11' of $\alpha$-tubulin | |
| 412-413, 416, 418-419, 421 | H12 of $\alpha$-tubulin | |
| 263-264 | Loop between H8 and B7 of $\beta$-tubulin | |
| 424, 428, 430, 432-435 | H12 of $\beta$-tubulin | |

[a] strong kinesin binding site identified in mutation experiment [1]

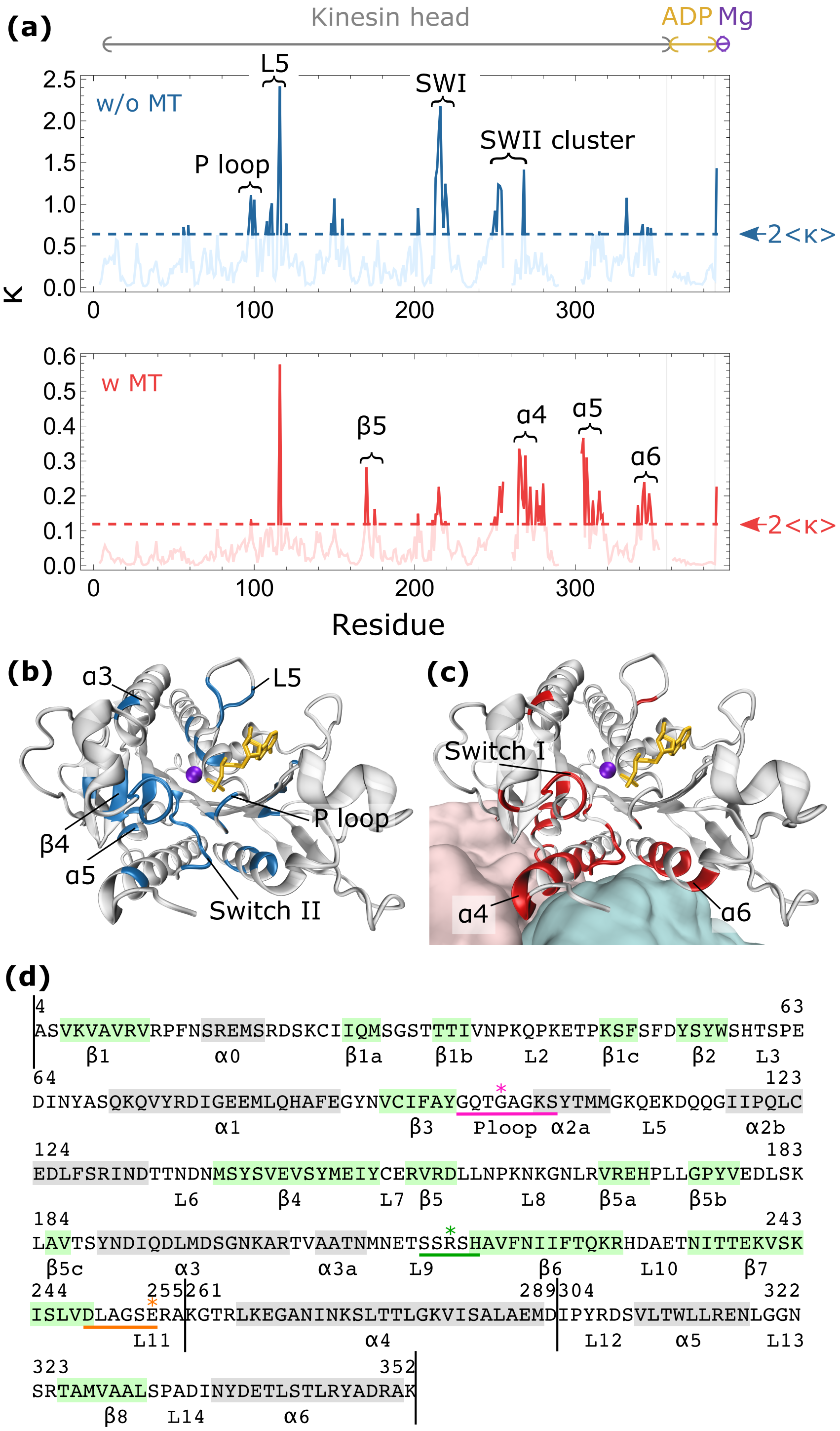


Figure S1: *See next page for caption*

Figure S1 *(previous page)*: Upon MT binding, residues in the MT binding regions of kinesin become stiffer. (a) To assess the relevance of a residue in transmitting allosteric signals, we calculate stiffness profiles of kinesin residues, $\kappa$ (Equation 3), for a monomeric kinesin kif1a (1i5s.pdb [2] – top panel) and a kif1a-MT complex (2hxh.pdb [3] – bottom panel), in the ADP bound state, using the 30 lowest energy normal modes. Residues that correspond to kinesin head (gray), atoms in ADP molecule (yellow), and $Mg^{2+}$ (purple) are marked on top and color-coded accordingly in Figure (b-c). Residues with $\kappa$ that are greater than twice the averaged value (dashed lines marked with arrow heads) are highlighted in Figure (b-c), listed in Table S1, and referred to as the Allosteric Wiring Diagram (AWD) [4]. (b) AWD in the kinesin structure (1i5s.pdb) are highlighted in blue. They include residues in switches I and II, P loop region, and $Mg^{2+}$. (c) Same as (b) except the AWD, in red, are for the kinesin head bound to the MT (2hxh.pdb). The $\alpha$ and $\beta$ structures in tubulin are presented in light cyan and pink. Besides the residues in switch I, II, P loop and $Mg^{2+}$, those near the MT binding sites ($\alpha 4$ and $\alpha 6$) have high values after MT binding. (d) Sequence of 1i5s.pdb with secondary structural elements (highlighted in silver for helices and green for strands). The vertical black lines are positions of the missing residues. Conserved motifs for P loop (magenta), switch I (green) and switch II (orange) are underlined.

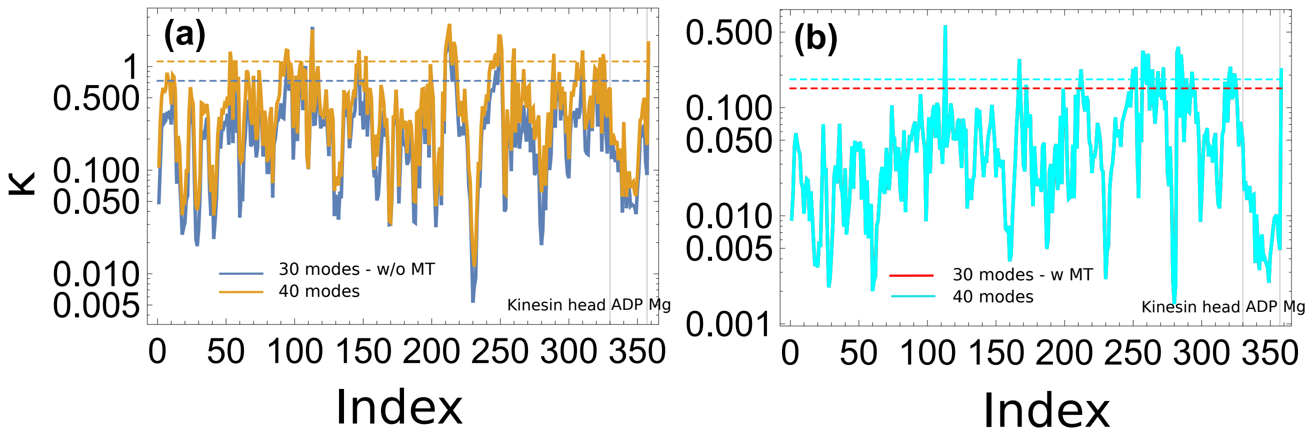


Figure S2: Comparisons between calculations with 30 modes vs 40 modes of Figure S1(a) for the data with (b) and without (a) MT. Increasing number of modes in the calculations does not significantly affect the main results.

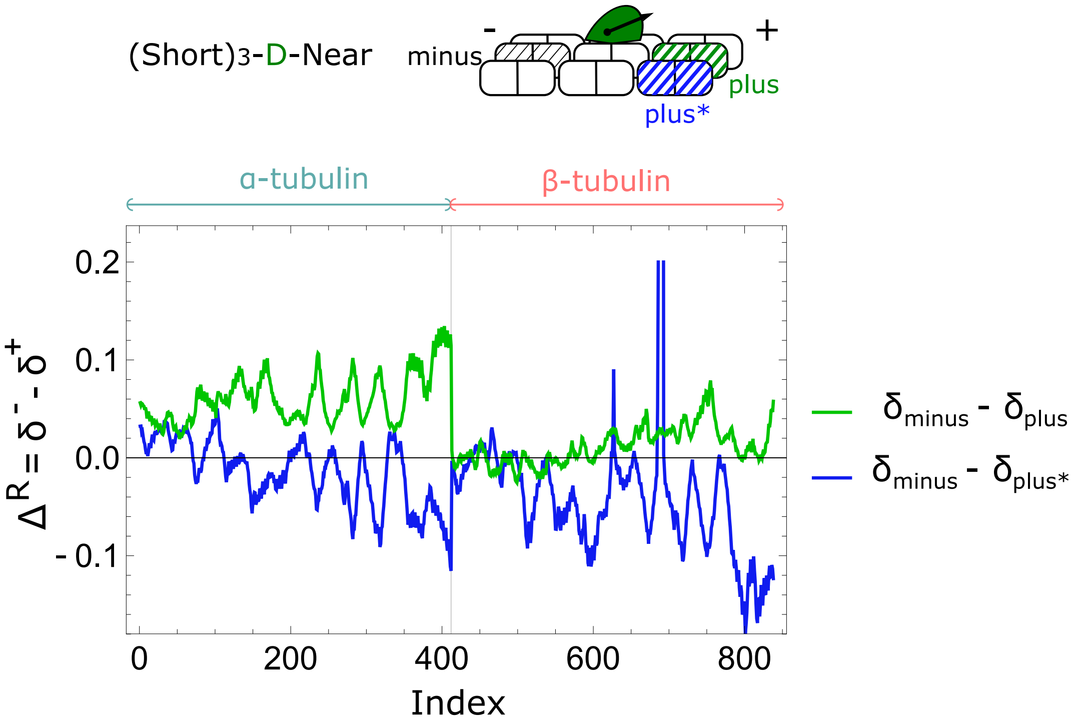


Figure S3: **Asymmetric response of MT upon binding of kinesin to the trailing head.** A MT structure fragment with 3 protofilaments, each with 3 $\alpha/\beta$ tubulins, was used in the calculations. The green line shows the difference $\delta_{minus} - \delta_{plus}$ due to the binding of the trailing head of kinesin, as in Figure 1(b) of the main text. The blue line represents the difference $\delta_{minus} - \delta_{plus^*}$ where $\delta_{plus^*}$ corresponds to a potential binding site for a side step.

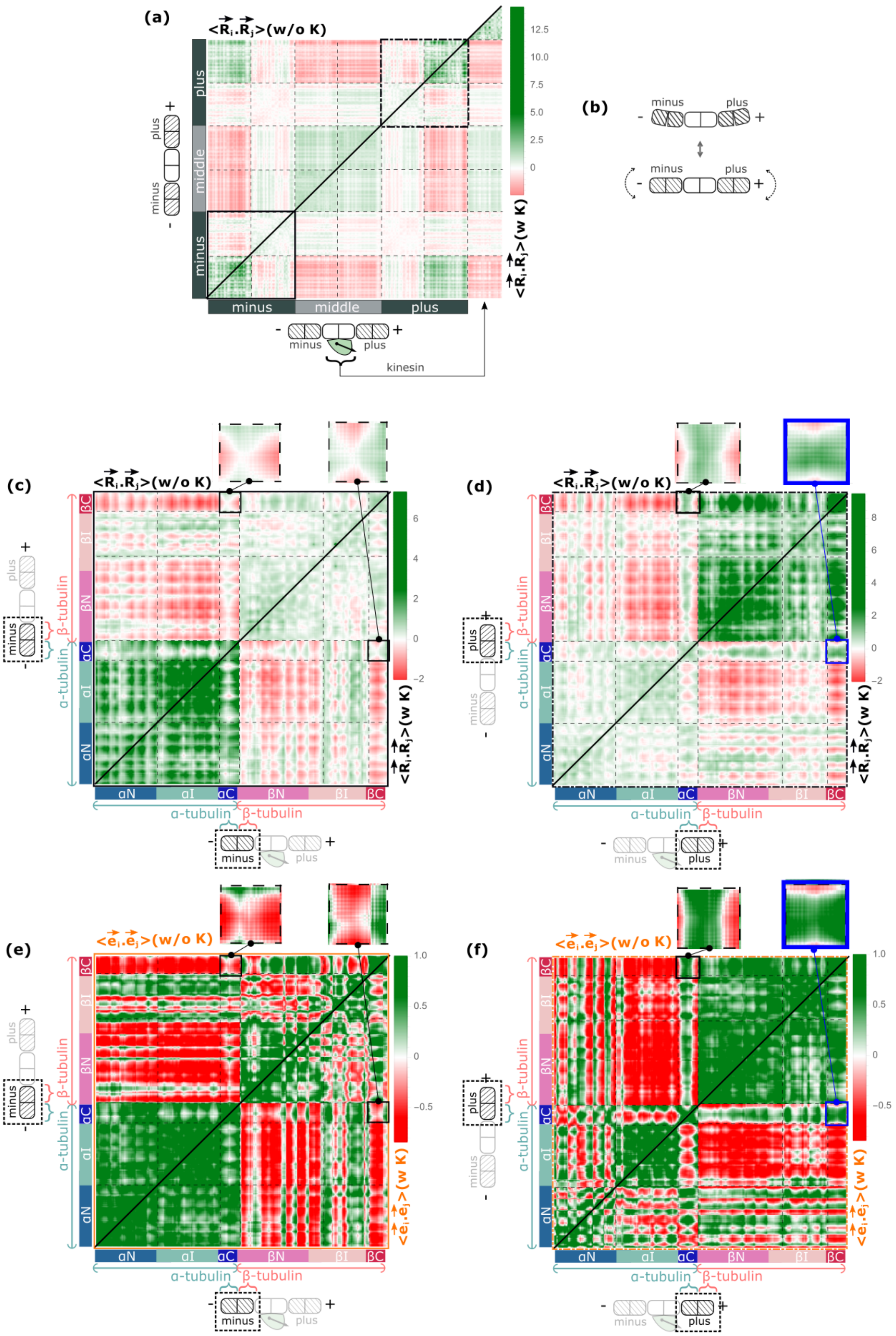


*See next page for caption*

Figure S4 *(previous page)*: C-terminal domains of the plus and minus end tubulins show the most significant asymmetric effect in the correlation profiles upon kinesin binding. (a) Dynamic correlation matrices $\langle \vec{R}_i \cdot \vec{R}_j \rangle$ between residues in a 3 tubulin dimer structure with and without bound kinesin. Positive and negative correlation values are shown in green and red hue, respectively. Both matrices show the most dominant mode: the bending mode as demonstrated in (b). (c-d) Dynamic correlation matrices $\langle \vec{R}_i \cdot \vec{R}_j \rangle$ and (e-f) orientational correlation matrices $\langle \vec{e}_i \cdot \vec{e}_j \rangle$ between residues in the minus (c, e) and plus (d, f) end tubulin dimers, with and without kinesin. The figures on top show the correlation matrices between the C-terminal domains $\alpha$C and $\beta$C.
